# Identification and classification of hubs in miRNA target gene networks in human neural stem/progenitor cells following Japanese encephalitis virus infection

**DOI:** 10.1101/581983

**Authors:** Sriparna Mukherjee, Irshad Akbar, Reshma Bhagat, Bibhabasu Hazra, Arindam Bhattacharyya, Pankaj Seth, Dipanjan Roy, Anirban Basu

**Author notes:** Correspondence: Anirban Basu and/or Dipanjan Roy Anirban Basu (Orcid id: http://orcid.org/0000-0002-5200-2054) Dipanjan Roy (Orcid id: https://orcid.org/0000-0002-1669-1083).

## Abstract

Micro RNA dysregulation is observed in many viral diseases. RNA viruses modulate host miRNA machinery for their own benefit. JEV, a neurotropic RNA virus has been reported to manipulate several miRNAs in neuron or microglia. However, no report indicates a complete sketch of the miRNA profile of NSPCs contributing to viral persistence; hence being focused in our current study. We performed a miRNA array of 84 miRNAs in human neuronal progenitor cell line and primary neural precursor cells isolated from aborted foetus. Several fold down-regulation of hsa-miR-9-5p, hsa-miR-22-3p, hsa-miR-124-3p and hsa-miR-132-3p were found in both of the cells. Subsequently, we screened for the target genes of these miRNAs and looked for the important biological pathways the genes significantly regulate. Then we sorted out the target genes which are involved in two or more than two pathways. We constructed a protein-protein interaction (PPI) network of the miRNA target genes based on their interaction patterns. A binary adjacency matrix for each gene network was prepared. Different modules or communities were identified in those networks using community detection algorithms. Mathematically, we identified the hub genes by analyzing their degree centrality and participation co-efficient in the network. The hub genes were classified either as provincial (P<0.4) or connector hubs (P>0.4). We validated the expression of hub genes in both cell line and primary cells through qRT-PCR post JEV infection and respective miR-mimic transfection. Taken together, our findings highlight the importance of specific target gene networks of miRNAs affected by JEV infection in NSPCs.

**Importance:** JEV damages the neural stem/progenitor cell population of mammalian brain. However, JEV induced alteration in miRNA expression pattern of the cell population remains an open question, hence warrants our present study. In this study, we specifically address the down-regulation of four miRNAs and we prepared a protein-protein interaction network of miRNA target genes. We identified two types of hub genes in the PPI network namely connector hubs and provincial hubs. These two types of miRNA target hub genes critically influence the participation strength in the networks and thereby significantly influence up and down regulation in several key biological pathways. Computational analysis of the PPI networks identifies key protein interactions and hubs in those modules which opens up the possibility of precise identification and classification of host factors for viral infection in NSPCs and how RNA viruses modulate host miRNA machinery for their own benefit post JEV infection to the cells.

## Introduction

Japanese encephalitis virus, transmitted by an arthropod vector, effectively replicates inside the central nervous system (CNS) and causes neuronal death. JEV also infects the mitotically active neural stem/progenitor cell population (NSPC) residing in the sub-ventricular zone and creates imbalance in endoplasmic reticulum homeostasis, thus activate cell death pathways^1,2^. JEV infection in NSPCs is linked to cognitive and motor deficiencies in the survivors. Thus, JEV promotes a double-trouble to brain by damaging the neuron and diminishing the stem-cell pool as well, so that CNS repair mechanism is hindered. miRNAs play a pivotal role in CNS development. Several miRNAs have been reported to regulate neurogenesis, neuronal migration, gliogenesis, synaptogenesis, long term potentiation and synaptic plasticity ^3^. They also regulate neural stem cell self-renewal and fate determination ^4^. Therefore, investigation of NSPC miRNA profile upon neurotropic virus invasion is of utmost importance.

The first miRNA was discovered in 1993 ^5^. Since then a radical research progress has happened in discovering their synthesis and biological importance. These 20-25nt long RNAs have been shown to bind to the 3′-UTR (un translated region) of target genes and thus restrain their translation ^6^.These small non-coding RNAs fine tune critical cellular processes in stem cells including cell cycle control, proliferation, differentiation and apoptosis ^7^. Aberrant expression of various miRNAs are documented in cases of Alzheimer’s disease^8^, Parkinson’s disease ^9^, Amyotrophic Lateral Sclerosis ^10^, Schizophrenia^11^ and Autism ^12^. miRNAs function as disease biomarkers and monitoring miRNA levels in a biological system provides a comprehensive picture of disease progression. Since miRNAs govern the expression of 50% protein coding genes in mammals, targeting them in a disease state is an appealing therapeutic strategy ^13^.

Tissue specific miRNA response upon viral challenge can directly alter viral replication and pathogenesis. Several studies report for a close connection of RNA virus infection and modulation of host miRNAs ^14,15^. As for example, Eastern equine encephalitis virus, (an alpha virus) infection modulates miR-142-3p expression in human and mouse macrophages ^16^ thus affecting viral propagation. Similarly, upon Picornavirus and Orthomyxovirus infections also modify certain host miRNAs ^17,18^. Neurotropic viruses can exploit the miRNA machinery in NSPCs. Down-regulation of miR-155 has been observed in HIV infection in neural precursor cells ^19^. Couple of other investigations also evaluates host miRNA reorganization following JEV infection. miR-15b has been shown to regulate inflammatory response in JEV infected microglial cells in a murine model ^20^. Alongside, miR-29b and miR-155 were also found to regulate JEV induced neuroinflammation^21,22^. A recent study indicates role of miR-301a inhibiting type I interferon signaling during JEV infection, thus regulating host immune response^23^. However, comprehension regarding miRNA response in NSPCs post JEV infection is still undiscovered.

Our group employed an in-silico tool to assess miRNA alteration following JEV infection in both hNS1 cells and neural precursor cells isolated from aborted human foetus. Through an analysis including target prediction, community detection, miRNA target hub genes identification and molecular validation, we illustrated miRNA-target gene interaction in case of JEV infection. These miRNAs might promote viral propagation and persistence in NSPCs, thus warrants further studies. We also anticipate that our study will open up the possibility of precise identification and classification of primary interaction partners which are key host factors for viral infection in NSPCs and how RNA viruses modulate host miRNA machinery for their own benefit post JEV pathogenesis in NSPCs.

## Materials and Methods

### Cell culture

#### hNS1 culture

hNS1 cells, an embryonic forebrain derived neural stem cell line, is a kind gift from Dr. Alberto Martínez-Serrano, Centre of Molecular Biology Severo Ochoa, Autonomous University of Madrid, Spain. hNS1 (formerly called HNSC.100, a model cell line of hNSCs) cells were grown according to previously used protocol ^1^. Briefly, cells were cultured in poly-D-lysine (Sigma) coated flasks in DMEM-F12 media (Invitrogen, CA, USA) containing 20ng/ml each of recombinant human EGF and FGF (R & D Systems), 1% BSA (Sigma), gentamycin and 1X N2 supplement (Invitrogen). Upon 65% confluency, cells were treated with 5 MOI of JEV and cells were harvested at 72 hrs post infection which has been previously studied to show significant viral infection ^1^.Mock infection is done by adding the same amount of media used for JEV infection but without virus.

Cell line authentication has been performed recently using multiplex STR system. The STR profiles verified were D5S818, D7S820, D16S539, TPOX, vWA, PentaE and Amelogenin.

#### Human neural precursor cells (hNPCs) culture

Isolation of hNPCs from aborted human foetus was carried out according to the ethical guidelines of NBRC, Department of Biotechnology (DBT) and Indian Council of Medical Research (ICMR) for Stem Cell Research. Briefly, cells were cultured in poly-D-lysine (Sigma) coated flasks in neurobasal media (Invitrogen) containing Neural Survival Factor (Lonza), N2 supplement (Invitrogen) 25 ng/ml bovine fibroblast growth factor (bFGF) (Sigma-Aldrich) and 20 ng/ml epidermal growth factor (EGF) (Sigma-Aldrich). Upon 65% confluency, cells were treated with 5 MOI of JEV and cells were harvested at 72 hrs post infection. Mock infection is done by adding the same amount of media used for JEV infection but without virus.

#### JEV infection in cells

Both hNS1 and hNP cells were infected with 5 MOI of JEV for 72 hrs and harvested for infection studies, miRNA isolation and RNA isolation. Control cells were treated with equal volume of PBS.

#### MicroRNA array

miRNA isolation from hNS1 cells and foetal neural stem cells were performed using miRNeasy mini kit (Qiagen, CA, USA) following manufacturer’s instruction. cDNA preparation was done using miScript II RT kit (Qiagen). The reaction condition was 37°C for 60 min and 95°C for 5 min. miScript miRNA PCR array of Human Neurological Development and Disease was performed using miScript SYBR green PCR kit (Qiagen). The PCR array contains 84 miRNAs which are differentially expressed during neuronal development and are responsible for progression of neurological diseases.

#### Real time PCR of miRNA

To check the expression of hsa-miR-9-5p, hsa-miR-22-3p, hsa-miR-124-3p and hsa-miR-132-3p, miRNA isolation was performed from harvested control and JEV infected hNS1 and hNP cells. cDNA was prepared using miScript II RT kit (Qiagen) as mentioned earlier. Primers for respective miRNA (Qiagen) were used in qRT-PCR reactions which are as follows:

miR-9-5p:5′UCUUUGGUUAUCUAGCUGUAUGA3′

miR-22-3p:5′AAGCUGCCAGUUGAAGAACUGU3′

miR-124-3p: 5′UAAGGCACGCGGUGAAUGCC3′

miR-132-3p: 5′UAACAGUCUACAGCCAUGGUGC3′

miScript SYBR green PCR kit (Qiagen) was used for all qRT-PCR reactions. The reaction condition for qRT-PCR was: 95°C for 15 min (1 cycle), 40 cycles at 94°C for 15 sec, 55°C for 30 sec and 70°C for 30 sec. SNORD68 was used as internal control. Data analysis was done using comparative delta CT method.

#### Transfection of cells with miRNA-mimic

To overexpress miR-9-5p, miR-22-3p, miR-124-3p and miR-132-3p, both hNS1 and hNP cells were transfected with human miRNA-mimics (double stranded RNAs that mimic mature endogenous miRNAs, Procured from Qiagen) of the mentioned. HiPerfect Transfection reagent (Qiagen) was used according to manufacturer’s protocol. 24 hrs post transfection cells were infected with JEV for specific time points. Controls of the mimic (Ambion) were used in all transfection experiments.

#### RNA isolation and real time PCR for target genes

hNS1 and hNP cells were harvested from culture plates using Trizol reagent (Sigma, USA) followed by chloroform treatment. These samples were centrifuged at 12000 rpm for 15 min at 4°C for phase separation. Aqueous phase was carefully collected and isopropanol was added to it followed by a centrifugation at 12000 rpm for 15 min at 4°C to get RNA pellet. RNA pellet was washed in 75% ethanol and air dried. cDNA was synthesized with 100 ng RNA using Advantage RT-PCR Kit (Clontech, Mountain View, USA). Reaction condition for real time PCR was 95°C for 3min (1 cycle), 95°C for 30 sec, annealing temperature for 30 sec and 72°C for 30 sec (45 cycles). The results were normalized using human GAPDH by ∆∆CT method and represented as fold change. Primer sequences are enlisted in Table S3.

#### MicroRNA isolation from human autopsy tissue

miRNA was isolated from paraffin embedded basal ganglia tissue of human autopsy cases validated to have JEV infection (CSF positive for IgM). Control tissue was collected from subjects died due to road accidents and do not have a prior record of infection in brain. The tissue samples were collected from Human Brain Bank, NIMHANS, Bangalore following institutional ethical rules. Deparaffinization was carried out according to our previously published article^24^. Then the tissue samples were homogenized using Qiazol reagent (Qiagen) and rest of the isolation procedure was performed using miRNeasy mini kit (Qiagen).

#### Graph theoretic analysis of miRNA target gene PPI networks

Graphs are composed of vertices (or nodes, here equivalent to target genes) and edges (or connections, here equivalent to inter-gene connectivity patterns). The connectivity structure of a graph is represented typically by its adjacency matrix; here we first construct an asymmetric binary matrix representing directed but unweighted edges. Paths are ordered sequences of edges linking pairs of vertices (a source and a target). The distance between two vertices corresponds to the length (number of edges) of the shortest path between them. The distance matrix of a graph comprises all pair-wise distances. Its maximum corresponds to the graph diameter, its minimum to the graph radius, and its average to the graph’s characteristic path length.

Basic graph measures such as connection density, proportion of reciprocal connections, degree distributions, measures derived from the distance matrix (diameter, radius, path length), and participation coefficients were calculated using standard graph theory methods. Brain Connectivity Toolbox (BCT) in MATLAB was used to compute modularity score and the participation coefficient of all the four miRNA target genes ^25^.

#### Community Structure Identification in target gene PPI networks

Modularity score is used to measure the community structure within a network. The value of modularity ranges between [−0.5, 1] with 0 and negative values meaning a network with randomly assigned edges to positive values indicating highly communal structure. In a given graph G (V, E) which can be partitioned into two membership variables *s*. If a node *i* falls into community 1 then *s*_*i*_ = 1 or else *s*_*i*_ = −1. An adjacency matrix may be denoted by *A*, which says *A*_*ij*_ = 1 means there is a connection between nodes *i* and *j* and *A*_*ij*_ = 0 when there are no interactions. Modularity (*Q*) is then defined as the fraction of edges that fall within community 1 or 2, minus the expected number of edges within communities 1 and 2 for a random graph with the identical node degree distribution as the given graph. All the modularity analysis is carried out by comparisons against populations of degree-matched random networks.

To identify modules (communities) within each target gene network, we apply a variant of a spectral community detection algorithm. Formally modularity (*Q*) can be defined as

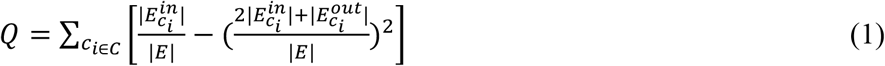

Where C is the set of all miRNA target gene communities, *c*_*i*_. is a specific community in C, 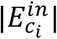 is the number of edges between node genes within community *c*_*i*_, 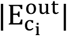 is the number of edges from genes in community *c*_*i*_ to target genes outside community *c*_*i*_, |E| is the total number of edges in the gene network.

For simplicity, modularity in Eq. (1) can also be expressed in a more compact form

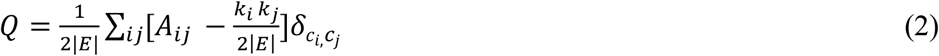

Where *k*_*i*_ is the degree of node i, *A*_*ij*_ is an element of the adjacency matrix, *δ*_*ci,cj*_ is the Kronecker delta symbol, *c*_i_ is the label of the community to which a target gene node *i* is being assigned. The modularity measure defined above is suitable only for undirected and unweighted networks. However, this definition can be naturally extended to apply to directed networks as well as to weighted networks. Weighted and directed networks contain more information than undirected and unweighted ones and are therefore often useful viewed as more valuable but also as more difficult to analyze than their simpler counterparts. The revised definition of modularity that works for directed networks is as follows

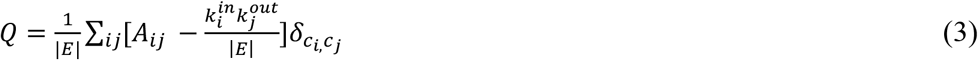

Where 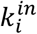 and 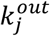 are the in- and out-degrees of the gene network. As inputs to the algorithm we used matrices of matching indices^25^, which express the similarity of connection patterns for each pair of vertices. Once modules were detected, different solutions were ranked according to a cost function and the optimal modularity (out of 10000 solutions for a range of between 2 and 6 modules) was used as the basis for hub classification. Details of the modularity optimization techniques used are outlined in the next subsection.

### Modularity Optimization of identified community structure

#### Spectral Methods and greedy search

Modularity in Eq. (3) can be re-written as

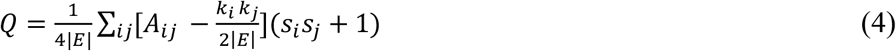

where *s* is column vector representing any division of the network into two groups. The elements of the column vector are defined as *s*_*i*_ = +1 if node *i* belongs to first group and *s*_*i*_ = −1 if the node belongs to the second group. The modularity matrix B with elements can be written as

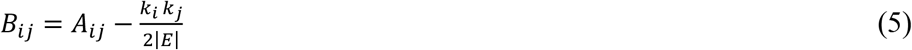

Representing *s* the column vector as a linear combination of the normalized eigenvectors ui of modularity matrix *B*: 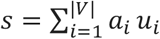 with 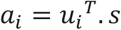, then plugging the result into Eq. (5) yield

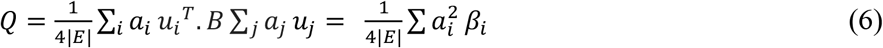

where *β*_*i*_ is the eigenvalue of B corresponding to eigenvector *u*_*i*_. To maximize Q above, Newman^26^ proposed a spectral approach to choose s proportional to the leading eigenvector *u*_*1*_ corresponding to the largest (most positive) eigenvalue *β*_*i*_. The algorithm that we used, initially divided the network into two communities and in further iterations the community structure is subdivided. The choice assumes that the eigenvalues are labeled in decreasing order *β*_*1*_ ≥ *β*_*2*_ ≥ … ≥ *β*_|*V*|_. Nodes are then divided into two communities according to the signs of the elements in s with nodes corresponding to positive elements in s assigned to one group and all remaining nodes to another. Since the row and column sums of B is zero, it always has an eigenvector (1, 1, 1, …) with eigenvalue zero. Therefore, if it has no positive eigenvalue, then the leading eigenvector is (1, 1, 1, …), which means that the network is indivisible. Once *Q* is almost 0 for an indivisible network, then further subdividing beyond this point will not contribute to the increase in modularity value *Q*. This can be used to terminate community structure division. Hence, the algorithm would end at a certain point when the optimal network has been estimated. In order to fine tune this method of community structure optimization further, this fitness function is typically performed using the Louvain method^26^, a greedy agglomerative clustering algorithm that works on hierarchical refinements of the network’s partitions. Here we used the Louvain implementation available in the Brain Connectivity toolbox^25^.

#### Classification of miRNA target Hub genes

To classify hub genes in the community which are potential miRNA targets we adopted the following strategy. First, we calculated each vertex’s participation index *P*, which expresses its distribution of intra-versus inter-module connections. *P* of vertex *v* is defined as ^27^

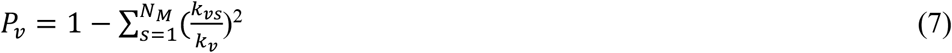

where *N*_*M*_ is the number of identified modules, *k*_*v*_ is the degree of node *v*, and *k*_*vs*_ is the number of edges from the *v*th node to nodes within module *s*. Considering only high-degree vertices (i.e. vertices with a degree at least one standard deviation above the network mean) we classify vertices with a participation coefficient *P*<0.4 as provincial hubs, and nodes with P>0.4 as connector hubs. Since *P* cannot exceed 0.5 for two-module networks and 0.67 for three-module networks, kinless hubs (i.e. nodes with P>0.8) cannot occur in these gene-gene interactome networks.

#### Statistical Analysis

Data is represented as mean±SD of three independent experiments (n=3). Statistical significance was calculated using Student’s *t*-test in case of two experimental groups or one-way analysis of variance (ANOVA) for multiple groups followed by Bonferroni *post hoc* test. p value < 0.05 was considered to be statistically significant.

## Results

### JEV infection alters miRNA expression profile of NSPCs

miRNA isolated from mock infected (control) and JEV infected hNS1 and hNPC samples were analyzed with Human Neurological Development and Disease miRNA PCR array in order to find out the differentially expressed miRNAs upon viral infection. Majority of the miRNAs were found to be down-regulated in both the cells upon infection (Fig 1). Color bars are indicative of fold change values in the infected cells compared to mock. Table S1 and S2 enlist the name and fold change value of the 84 miRNAs present in the mentioned array. Among the downregulated miRNAs we sorted out the common ones those are downregulated in both the cells and thus hsa-miR-9-5p, hsa-miR-22-3p, hsa-mir-124-3p and hsa-miR-132-3p were chosen for further experiments.

**Figure 1.**
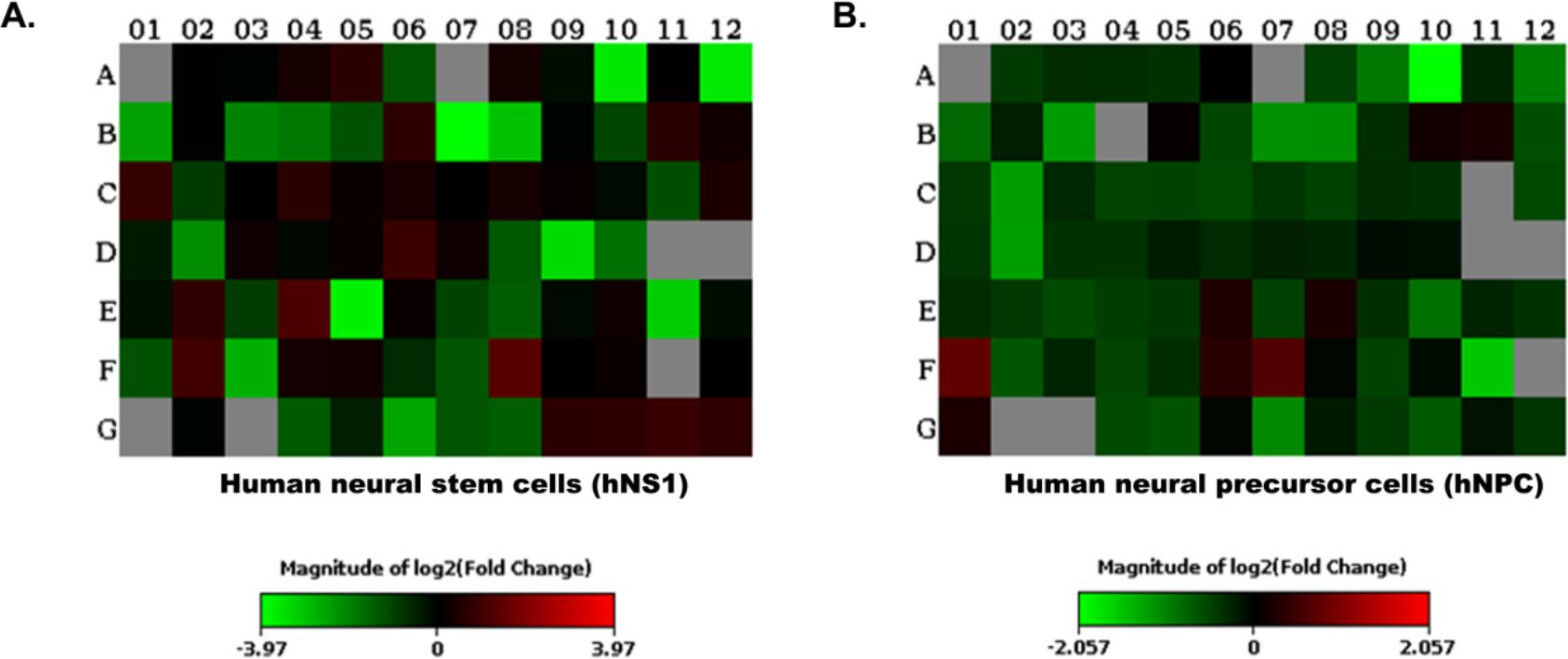
Heat map depicting differential expression of miRNAs in hNS1 and hNPC. miRNA was extracted from mock infected (control) and JEV infected hNS1 and hNPC samples and analyzed with Human Neurological Development and Disease miRNA PCR array. (A) Heat map showing differential expression of miRNAs in JEV infected hNS1 cells compared to mock. (B) Heat map showing differential expression of miRNAs in JEV infected hNP cells compared to control. Analysis was performed using: http://www.sabiosciences.com/pcrarraydataanalysis.php. Data is representative of three independent experiments. Color bars indicate the fold change value (green: down-regulation, red: up-regulation).

### Network analysis of hsa-miR-9-5p target gene networks and hub identification

In order to identify hsa-miR-9-5p target genes we used STRING 9.1 online database (http://string-db.org/) for extraction of the gene networks. These gene names and their repetition number in various pathways were listed in Table S4. There were about 60 genes identified by the string database. A comprehensive human gene-gene network constructed by defining an adjacency matrix *A*_*ij*_ was used for the network analysis. This adjacency matrix *A*_*ij*_ comprises of 60 target genes (nodes) and binarized edges based on their functional annotations (Fig 2A, B). Subsequently, modularity score *Q* were computed using Brain Connectivity Toolbox (BCT) in MATLAB and genes were partitioned to six identified communities based on similarity of connection patterns for each pair of vertices/nodes based on Eq. (1,2, and 3) and plotted in Fig 2C. This process of community detection was repeated using greedy search procedure based on Louvain method as displayed in Eq. (4, 5 and 6). In order to visualize these six communities and their interactomes Cytoscape (https://cytoscape.org/) an open source online platform was used to study community level specific interaction patterns. The network modularity analysis further confirmed there were several dominant players in biological pathway underlying JEV infection in neurons. In order to find out which genes may serve as hub genes and modulate more than one subnetwork which were identified using modularity analysis we have employed a hub identification algorithm. Based on the node degree *k*_*v*_ of node *v*, and *k*_*vs*_ is the number of edges from the *v*th node to nodes within module *s*, we have estimated each vertex’s participation index *P* and quantified the presence of provincial and connector hub genes in the interactome network. The results of hub analysis were displayed in Fig 2D. Our hub analysis predicted there were a number of connector hub genes present in each module including SIRT1, PTGS2, ETS1, SUMO1, IL6 etc. which could possibly have key role in up and down regulation in key biological pathways.

**Figure 2.**
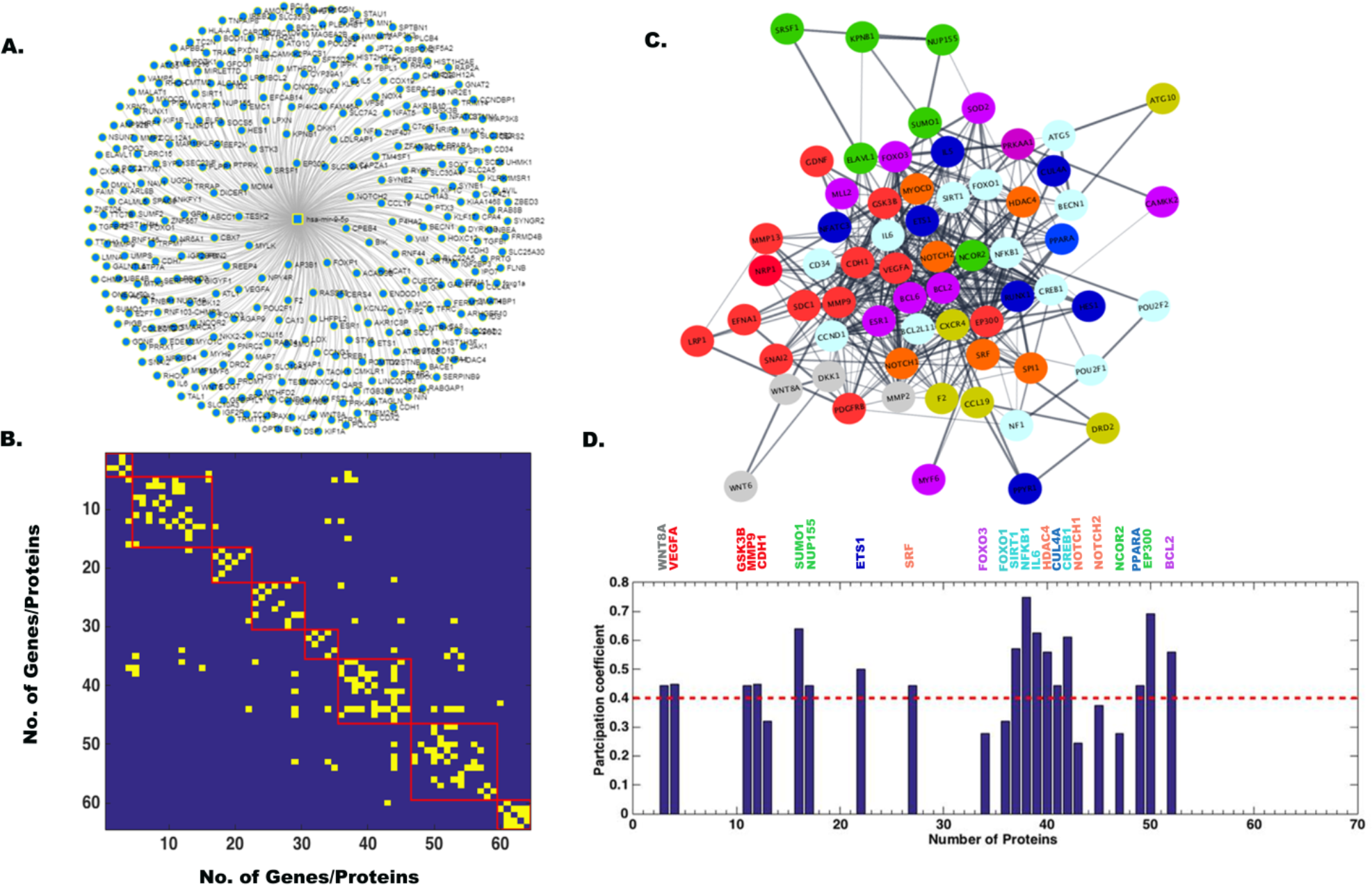
Network analysis showing miR-9-5p target gene networks, modularity/community detection in gene networks and the identification of hub genes based on participation coefficients. There were about 60 genes identified by the string database (A). A comprehensive human gene-gene network constructed by defining an adjacency matrix A_ij_ *A*_*ij*_ was used for the network analysis. This adjacency matrix *A*_*ij*_ comprises of 60 target genes (nodes) and binarized edges based on their functional annotations (B). Modularity score *Q* were computed using Brain Connectivity Toolbox (BCT) in MATLAB and genes were partitioned to eight identified communities based on similarity of connection patterns for each pair of vertices/nodes (C). Based on the node degree *k*_*v*_ of node *v*, and *k*_*vs*_ is the number of edges from the *v*th node to nodes within module *s*, we have estimated each vertex’s participation index *P* and quantified the presence of provincial and connector hub genes (D). The participation index *P* shows that there are provincial and connector hub genes belonging to seven communities.

### Network analysis of hsa-miR-22-3p target gene networks and hub identification

In order to identify hsa-miR-22-3p target genes as described in the previous section we used STRING 9.1 online database (http://string-db.org/) for extraction of the gene networks. All the identified genes and their repetition number in various biological pathways were listed in Table S5. There were about 40 genes identified by the string database displayed in Fig 3A. Interactomes were constructed by defining an adjacency matrix *A*_*ij*_. This adjacency matrix *A*_*ij*_ comprises of 40 target genes (nodes) and binarized edges between them plotted in Fig 3B. Subsequently, modularity score *Q* had been estimated and genes were partitioned to seven community structures based on similarity of connection patterns for each pair of vertices/nodes based on Eq. (1,2, and 3) and plotted in Fig 3C. This process of community detection was repeated using greedy search procedure as described in Eq. (4, 5 and 6). This network modularity analysis confirmed there were several dominant players in biological pathway underlying JEV infection in neurons. In order to find out which genes may serve as hub genes and modulate more than one subnetwork we have applied a hub detection algorithm. Based on the node degree *k*_*v*_ of node *v*, and *k*_*vs*_ is the number of edges from the *v*th node to nodes within module *s*, we have estimated each vertex’s participation index *P* and quantified the presence of provincial and connector hub genes in the interactome network plotted in Fig 3D. Our hub analysis predicted there were a number of connector hub genes e.g. CDKN1A, SIRT1, NR3C1, MAX1, NCOA1, ESR1, SP1 and PTEN etc. specifically targeted by the hsa-miR-22-3p. Next, we carried out a molecular validation whether any of the identified hub genes got up or down regulated post JEV infection in different cells.

**Figure 3.**
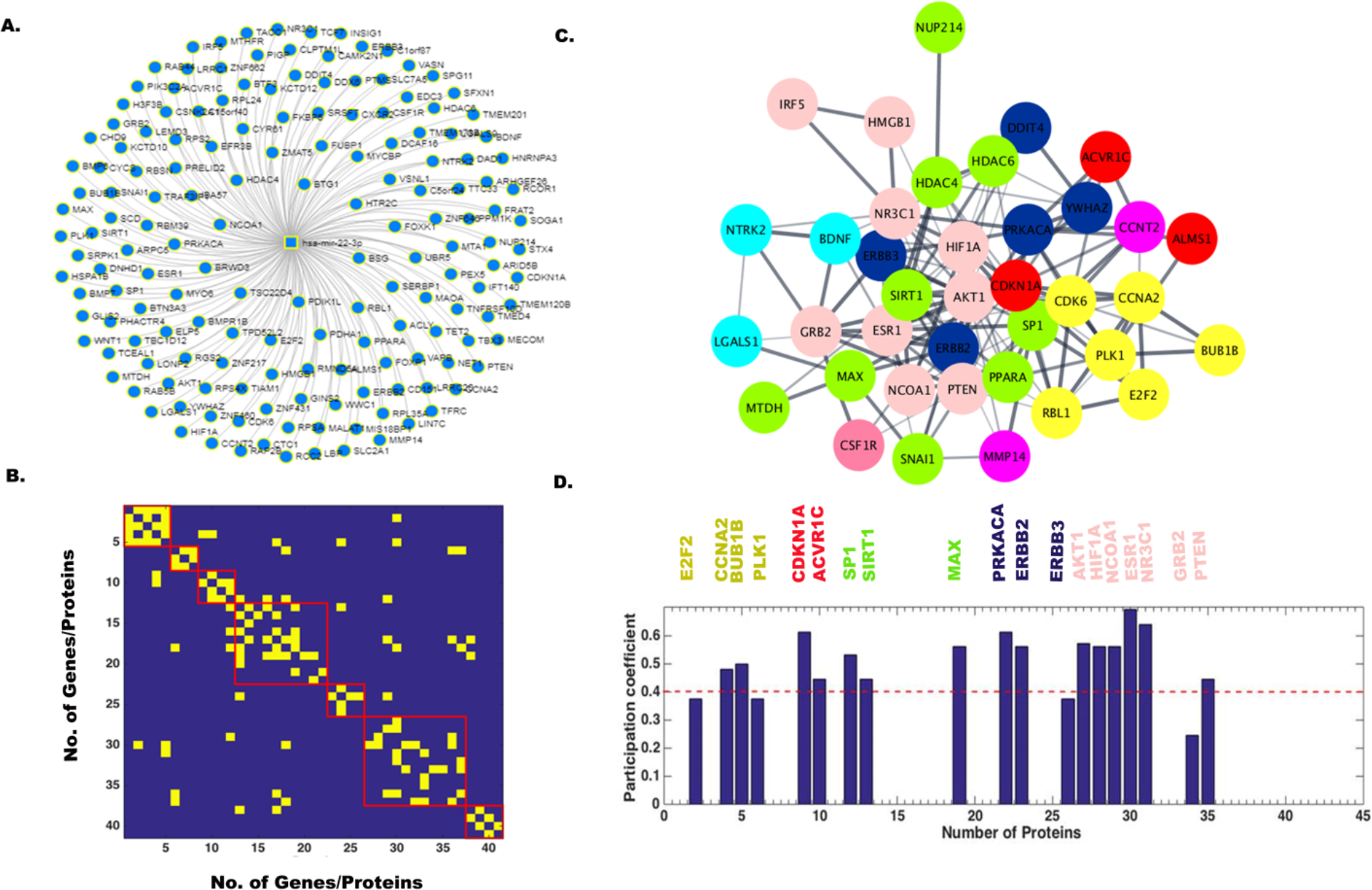
Network analysis showing miR-22-3p target gene networks, modularity/community detection in gene networks and the identification of hub genes based on participation coefficients. There were about 40 genes identified by the string database (A). A comprehensive human gene-gene network constructed by defining an adjacency matrix *A*_*ij*_ was used for the network analysis. This adjacency matrix *A*_*ij*_ comprises of 40 target genes (nodes) and binarized edges based on their functional annotations (B). Modularity score *Q* were computed using Brain Connectivity Toolbox (BCT) in MATLAB and genes were partitioned to seven identified communities based on similarity of connection patterns for each pair of vertices/nodes (C). Based on the node degree *k*_*v*_ of node *v*, and *k*_*vs*_ is the number of edges from the *v*th node to nodes within module *s*, we have estimated each vertex’s participation index *P* and quantified the presence of provincial and connector hub genes (D). The participation index *P* shows that there are provincial and connector hub genes belonging to five communities.

### Network analysis of hsa-miR-124-3p target gene networks and hub identification

In order to identify hsa-miR-124-3p target genes as described in the previous section we used STRING 9.1 online database (http://string-db.org/) for extraction of the gene networks. All the identified genes and their repetition number in various biological pathways were listed in Table S6. There were about 90 genes identified by the string database plotted in Fig 4A. Interactomes were constructed by defining an adjacency matrix *A*_*ij*_. This adjacency matrix *A*_*ij*_ comprises of 90 target genes (nodes) and binarized edges between them plotted in Fig 4B. Subsequently, modularity score *Q* had been estimated and genes were partitioned to five community structures based on similarity of connection patterns for each pair of vertices/nodes based on Eq. (1,2, and 3) as plotted in Fig 4C. This process of community detection was repeated using greedy search procedure as described in Eq. (4, 5 and 6). This network modularity analysis confirmed there were several dominant players in biological pathway underlying JEV infection in cells. In order to find out which genes in the respective communities may serve as hub genes and interact with more than one subnetwork we have applied a hub detection algorithm. Based on the node degree *k*_*v*_ of node *v*, and *k*_*vs*_ is the number of edges from the *v*th node to nodes within module *s*, we have estimated each vertex’s participation index *P* and quantified the presence of provincial and connector hub genes in the interactome network plotted in Fig 4D. Our hub analysis predicted there were a number of connector hub genes e.g. SDC4, PIK3CA, CAV1, PIM1, IQGAP1, NRP1, SHC1, MAP3K3, etc. specifically targeted by the hsa-miR-124-3p. Next, we carried out a molecular validation whether any of the identified hub genes got up or down regulated post JEV infection in different cells.

**Figure 4.**
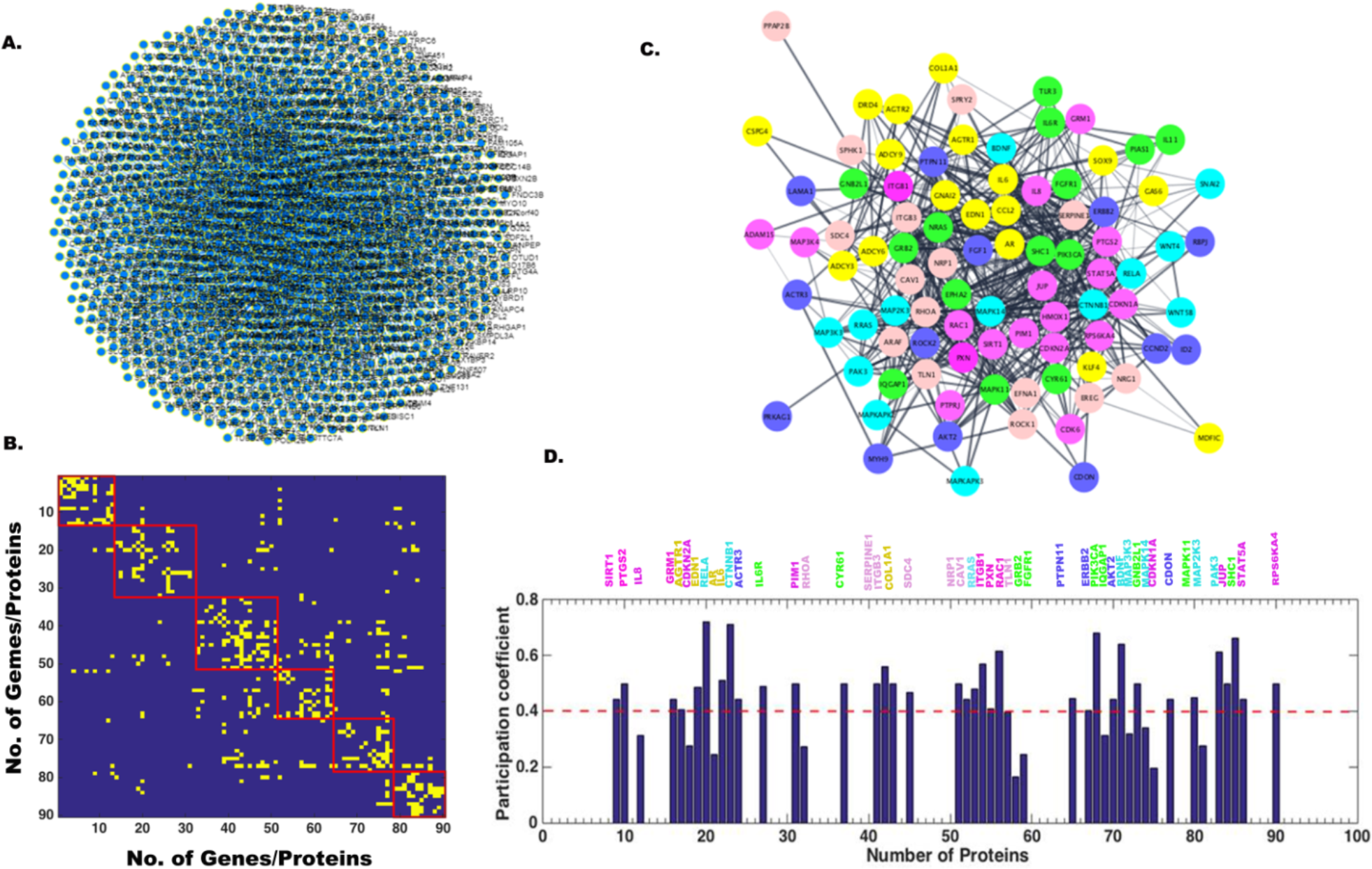
Network analysis showing miR-124-3p target gene networks, modularity/community detection in gene networks and the identification of hub genes based on participation coefficients. There were about 90 genes identified by the string database (A). A comprehensive human gene-gene network constructed by defining an adjacency matrix *A*_*ij*_ was used for the network analysis. This adjacency matrix *A*_*ij*_ comprises of 90 target genes (nodes) and binarized edges based on their functional annotations (B). Modularity score *Q* were computed using Brain Connectivity Toolbox (BCT) in MATLAB and genes were partitioned to six identified communities based on similarity of connection patterns for each pair of vertices/nodes (C). Based on the node degree *k*_*v*_ of node *v*, and *k*_*vs*_ is the number of edges from the *v*th node to nodes within module *s*, we have estimated each vertex’s participation index *P* and quantified the presence of provincial and connector hub genes (D). The participation index *P* shows that there are provincial and connector hub genes belonging to all the identified six communities.

### Network analysis of hsa-miR-132-3p target gene networks and hub identification

In order to identify hsa-miR-132-3p target genes as described in the previous section we used STRING 9.1 online database (http://string-db.org/) for extraction of the gene networks. All the identified genes and their repetition number in various biological pathways were listed in Table S7. There were about 20 genes identified by the string database plotted in Fig 5A. Interactomes were constructed by defining an adjacency matrix *A*_*ij*_. This adjacency matrix *A*_*ij*_ comprises of 20 target genes (nodes) and binarized edges between them plotted in Fig 5B. Subsequently, modularity score *Q* had been estimated and genes were partitioned to four community structures based on similarity of connection patterns for each pair of vertices/nodes based on Eq. (1,2, and 3) as plotted in Fig 5C. The process of community detection was repeated using greedy search procedure as described in Eq. (4, 5 and 6). This network modularity analysis further confirmed there were several dominant players in biological pathway underlying JEV infection in neurons. In order to find out which genes may serve as hub genes and modulate more than one subnetwork we have applied a hub detection algorithm. Based on the node degree *k*_*v*_ of node *v*, and *k*_*vs*_ is the number of edges from the *v*th node to nodes within module *s*, we have estimated each vertex’s participation index *P* and quantified the presence of provincial and connector hub genes in the interactome network plotted in Fig 5D. Our hub analysis predicted there were a number of connector hub genes e.g., RAF1, Rb1, MAPK1, RASA1, EGFR1, USP8 etc. specifically targeted by the hsa-miR-132-3p. Next, we carried out a molecular validation whether any of the identified hub genes got up or down regulated post JEV infection in different cells.

**Figure 5.**
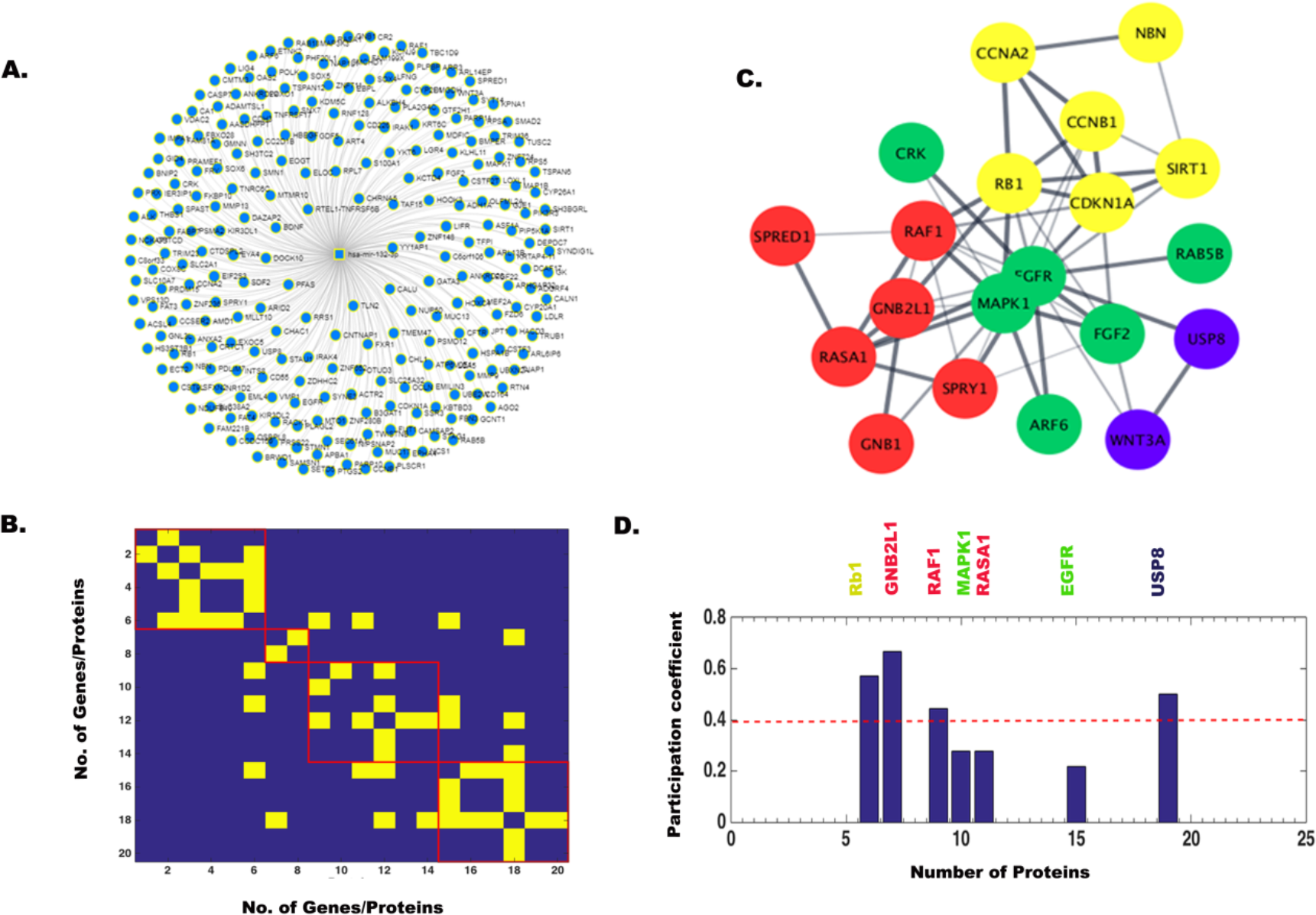
Network analysis showing miR-132-3p target gene networks, modularity/community detection in gene networks and the identification of hub genes based on participation coefficients. There were about 20 genes identified by the string database (A). A comprehensive human gene-gene network constructed by defining an adjacency matrix *A*_*ij*_ was used for the network analysis. This adjacency matrix *A*_*ij*_ comprises of 20 target genes (nodes) and binarized edges based on their functional annotations (B). Modularity score *Q* were computed using Brain Connectivity Toolbox (BCT) in MATLAB and genes were partitioned to four identified communities based on similarity of connection patterns for each pair of vertices/nodes (C). Based on the node degree *k*_*v*_ of node *v*, and *k*_*vs*_ is the number of edges from the *v*th node to nodes within module *s*, we have estimated each vertex’s participation index *P* and quantified the presence of provincial and connector hub genes (D). The participation index *P* shows that there are provincial and connector hub genes belonging to all the identified four communities.

### Expression of hsa-miR-9-5p target genes in hNS1 and hNP cells

hNS1 and hNP cells were divided into 5 experimental groups. One group was transfected with control mimic and two other groups with miR-9-5p mimic. 24 hrs post transfection, 3 groups were infected with 5 MOI of JEV (JEV, control mimic+ JEV and miR-9-5p mimic+JEV). Control group was treated with equal volume of PBS. 48 hrs post infection cells were harvested for RNA isolation and cDNA preparation. Expression of miR-9-5p target genes were validated through qRT-PCR with both hNS1 (Fig 6A) and hNP cell (Fig 6B) samples. Although ETS1, SIRT1, SUMO1 and IL-6 had up-regulation post JEV infection, miR-9-5p mimic transfection could not down-regulate all of their expressions. ETS1 and IL-6 were found to be significantly down-regulated upon mimic transfection in hNS1 cells, whereas only ETS1 was notably down-regulated in hNP cells on same experimental condition. GAPDH gene expression was taken as normalization control.

**Figure 6.**
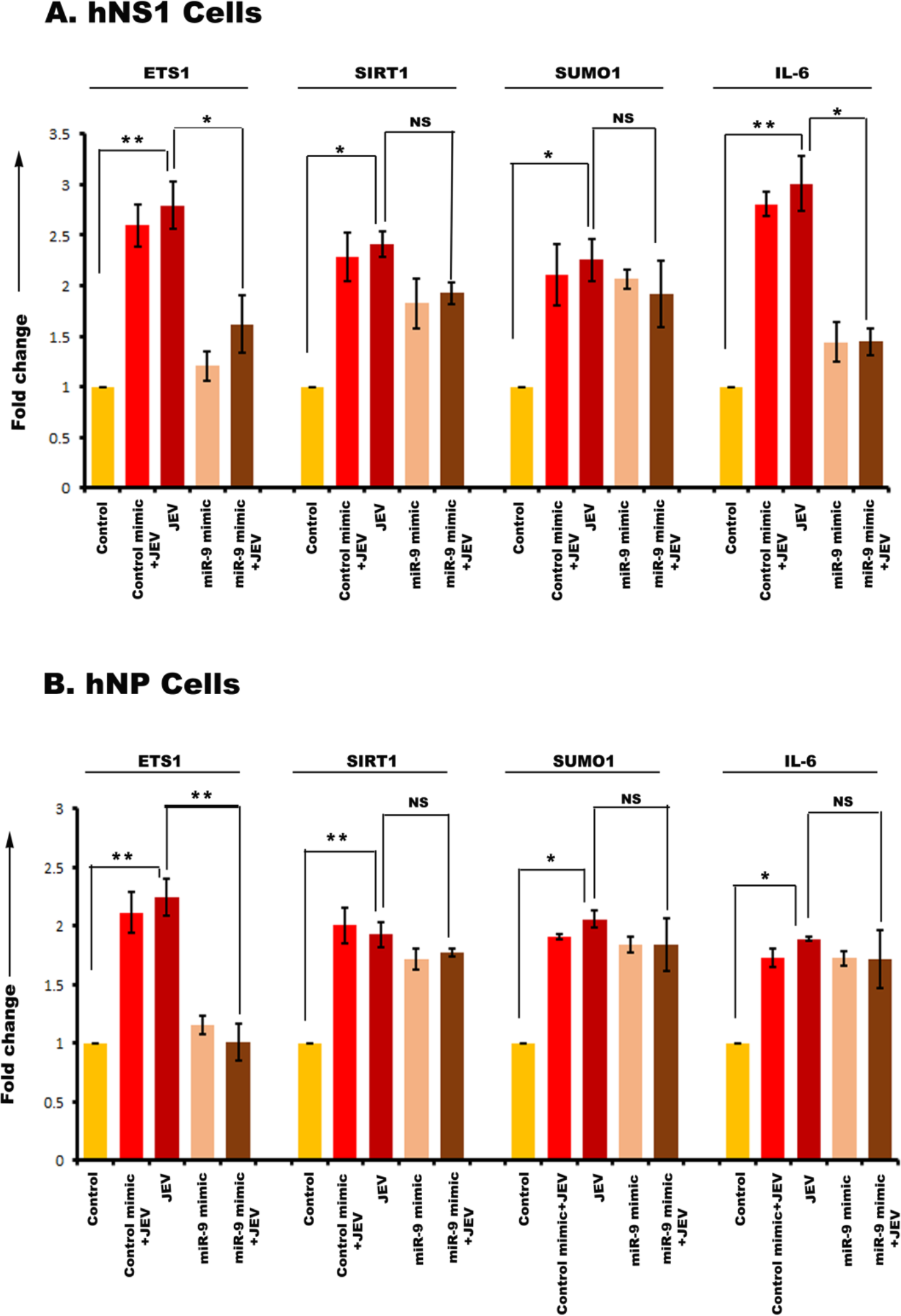
qRT-PCR showing miR-9-5p target gene expression in hNS1 and hNP cells upon JEV infection and miR-mimic transfection. RNA isolated from control, control mimic transfected+JEV infected, JEV infected, miR-9-5p mimic transfected and miR-9-5p mimic transfected+JEV infected hNS1and hNP cells were subjected to qRT-PCR. JEV infection led to significant up-regulation of ETS1, SIRT1, SUMO1 and IL-6 in both the cells (A, B). But upon mimic transfection and JEV infection, ETS1 and IL-6 genes showed a reduced expression profile in hNS1 cells (A). Whereas, in case of hNP cells, only ESR1 gene expression was down-regulated after mimic transfection and viral infection (B). GAPDH gene expression was used as an internal control for normalization of PCR data in both the cells. Data is representative of three independent experiments (mean ± SD) by one way ANOVA (**p<0.05, **p<0.01).

### Expression of hsa-miR-22-3p target genes in hNS1 and hNP cells

hNS1 and hNP cells were divided into 5 experimental groups. One group was transfected with control mimic and two other groups with miR-22-3p mimic. 24 hrs post transfection, 3 groups were infected with 5 MOI of JEV (JEV, control mimic+ JEV and miR-22-3p mimic+JEV). Control group was treated with equal volume of PBS. 48 hrs post infection cells were harvested for RNA isolation and cDNA preparation. Expression of miR-22-3p target genes were validated through qRT-PCR with both hNS1 (Fig 7A) and hNP cell (Fig 7B) samples. The target genes MAX1, NCOA1, NR3C1, ESR1, SP1 and PTEN had an up-regulation post JEV infection. Upon miR-22-3p mimic transfection; MAX1, NCOA1 and NR3C1 were found to be significantly down-regulated in hNS1 cells, whereas only ESR1 was notably down-regulated in hNP cells. GAPDH gene expression was taken as normalization control.

**Figure 7.**
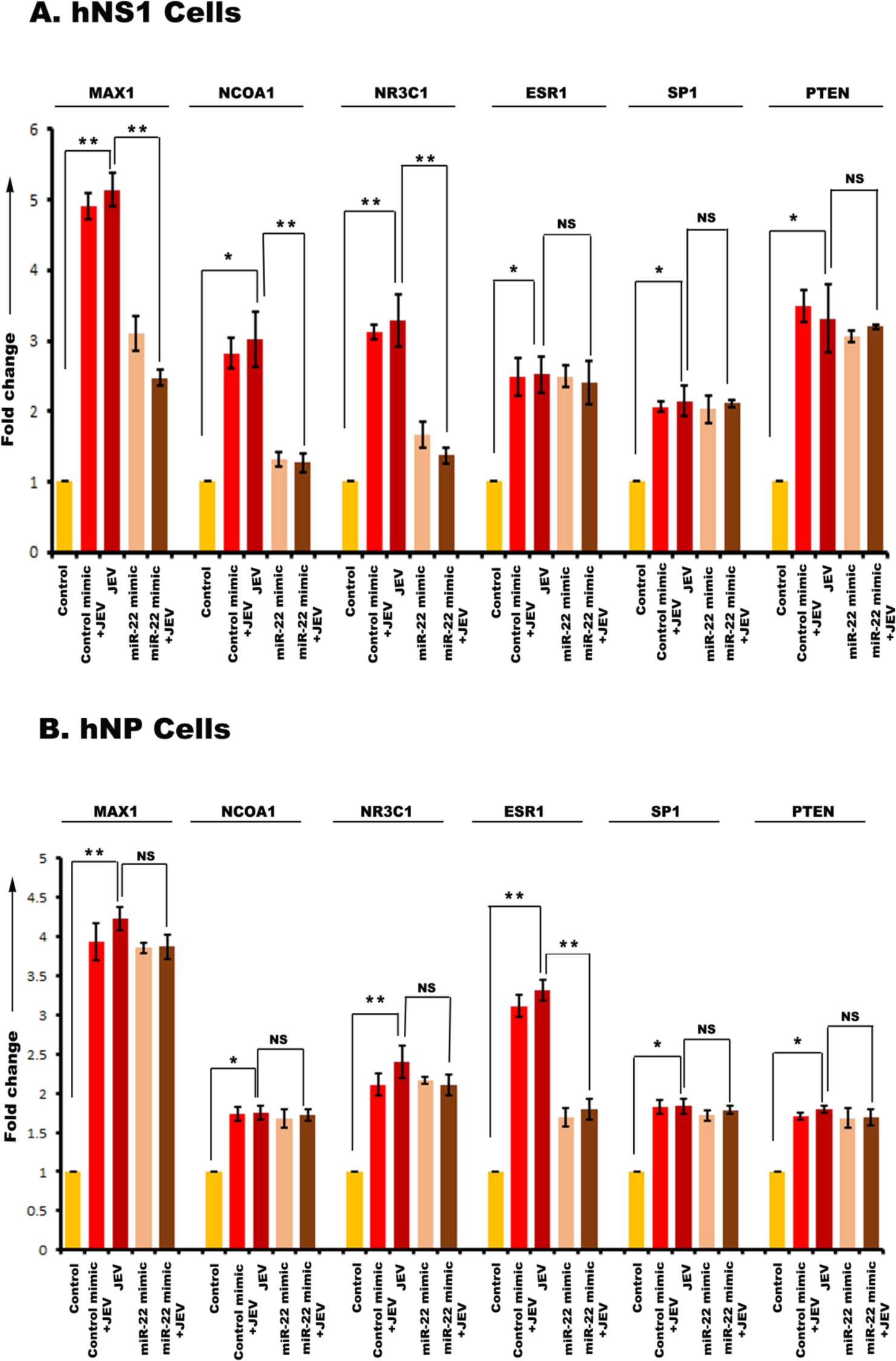
qRT-PCR showing miR-22-3p target gene expression in hNS1 and hNP cells upon JEV infection and miR-mimic transfection. RNA isolated from control, control mimic transfected+JEV infected, JEV infected, miR-22-3p mimic transfected and miR-22-3p mimic transfected+JEV infected hNS1and hNP cells were subjected to qRT-PCR. JEV infection led to significant up-regulation of MAX1, NCOA1, NR3C1, ESR1, SP1 and PTEN in both the cells (A, B). But upon mimic transfection and JEV infection, MAX1, NCOA1, NR3C1 genes showed a reduced expression profile in hNS1 cells (A). Whereas, in case of hNP cells, only ESR1 gene expression was down-regulated after mimic transfection and viral infection (B). GAPDH gene expression was used as an internal control for normalization of PCR data in both the cells. Data is representative of three independent experiments (mean± SD) by one way ANOVA (**p<0.05, **p<0.01).

### Expression of hsa-miR-124-3p target genes in hNS1 cells

hNS1 cells were divided into 5 experimental groups. One group was transfected with control mimic and two other groups with miR-124-3p mimic. 24 hrs post transfection, 3 groups were infected with 5 MOI of JEV (JEV, control mimic+ JEV and miR-124-3p mimic+JEV). Control group was treated with equal volume of PBS. 48 hrs post infection cells were harvested for RNA isolation and cDNA preparation. Expression of miR-124-3p target genes were validated through qRT-PCR with hNS1 cells (Fig 8). SDC4, PIK3CA, CAV1 genes showed up-regulated expression upon viral infection and after mimic transfection and infection, their expressions were decreased. Other genes (PIM1, IQGAP1, NRP1, SHC1, and MAP3K3) had an up-regulated profile post infection but mimic transfection had no effect on their expression. GAPDH gene expression was taken as normalization control.

**Figure 8.**
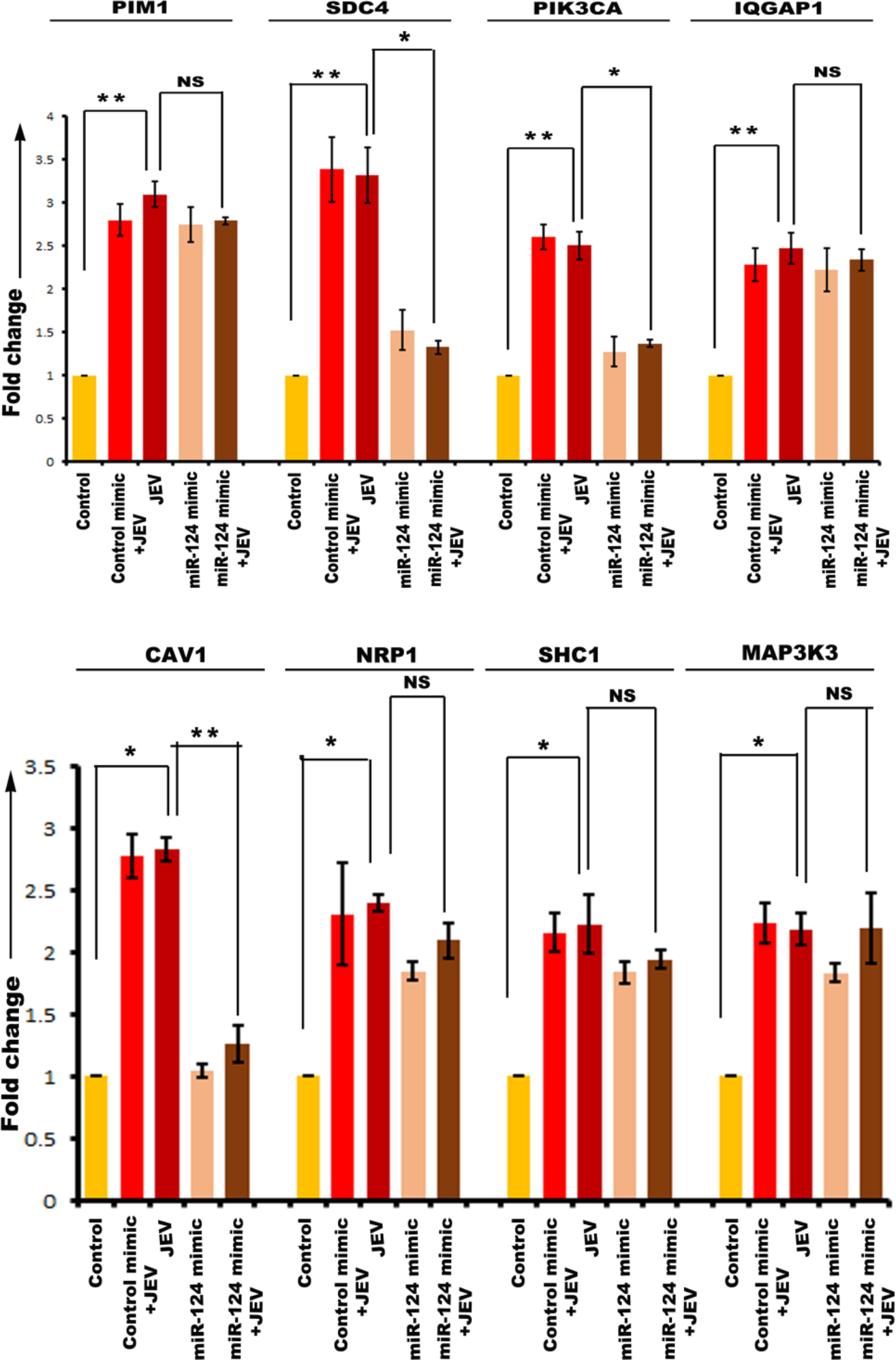
qRT-PCR showing miR-124-3p target gene expression in hNS1 cells upon JEV infection and miR-mimic transfection. RNA isolated from control, control mimic transfected+JEV infected, JEV infected, miR-124-3p mimic transfected and miR-124-3p mimic transfected+JEV infected hNS1cells were subjected to qRT-PCR. JEV infection led to significant up-regulation of PIM1, SDC4, PIK3CA, IQGAP1 (upper panel), CAV1, NRP1, SHC1 and MAP3K3 (lower panel). But upon mimic transfection and JEV infection, SDC4, PIK3CA and CAV1 genes showed a reduced expression profile. GAPDH gene expression was used as an internal control for normalization of PCR data. Data is representative of three independent experiments (mean± SD) by one way ANOVA (**p<0.05, **p<0.01).

### Expression of hsa-miR-124-3p target genes in hNP cells

hNP cells were divided into 5 experimental groups. One group was transfected with control mimic and two other groups with miR-124-3p mimic. 24 hrs post transfection, 3 groups were infected with 5 MOI of JEV (JEV, control mimic+ JEV and miR-124-3p mimic+JEV). Control group was treated with equal volume of PBS. 48 hrs post infection cells were harvested for RNA isolation and cDNA preparation. Expression of miR-124-3p target genes were validated through qRT-PCR with hNP cells (Fig 9). SDC4, PIK3CA, IQGAP1 genes showed up-regulated expression upon viral infection and after mimic transfection and infection, their expressions were decreased. Other genes (PIM1, CAV1, NRP1, SHC1, and MAP3K3) had an up-regulated profile post infection but mimic transfection had no effect on their expression. GAPDH gene expression was taken as normalization control.

**Figure 9.**
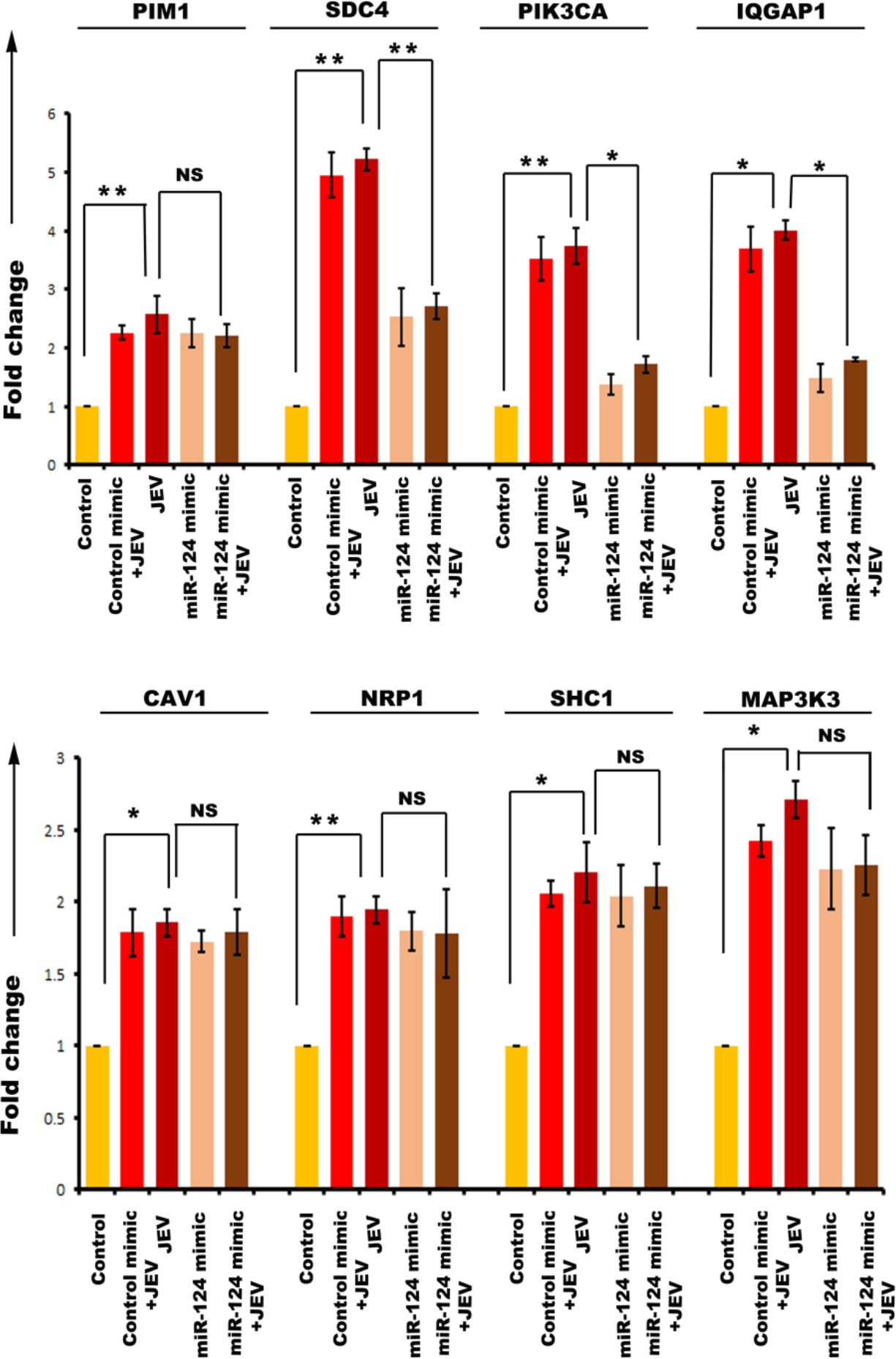
qRT-PCR showing miR-124-3p target gene expression in hNP cells upon JEV infection and miR-mimic transfection. RNA isolated from control, control mimic transfected+JEV infected, JEV infected, miR-124-3p mimic transfected and miR-124-3p mimic transfected+JEV infected hNP cells were subjected to qRT-PCR. JEV infection led to significant up-regulation of PIM1, SDC4, PIK3CA, IQGAP1 (upper panel), CAV1, NRP1, SHC1 and MAP3K3 (lower panel). But upon mimic transfection and JEV infection, SDC4, PIK3CA and IQGAP1 genes showed a reduced expression profile. GAPDH gene expression was used as an internal control for normalization of PCR data. Data is representative of three independent experiments (mean± SD) by one way ANOVA (**p<0.05, **p<0.01).

### Altered expression of miRNAs in human autopsy tissue of JEV infection cases

Expression of miR-9-5p, miR-22-3p, mir-124-3p and miR-132-3p were assessed in JEV infected and un-infected (non-JE) human autopsy samples. Significant reduction in expression of miR-9-5p and miR-22-3p were found in JEV infected samples compared to non-JE (Fig 10). Expressions of other two miRNAs remain unaltered (data not shown). This experiment was performed to get a parallel data of miRNA profile in human JE cases. Due to the unavailability of JEV infected neurogenic region (SVZ) in NIMHANS brain repository, this experiment was done with basal ganglia tissue. This is an impending limitation to our present study.

**Figure 10.**
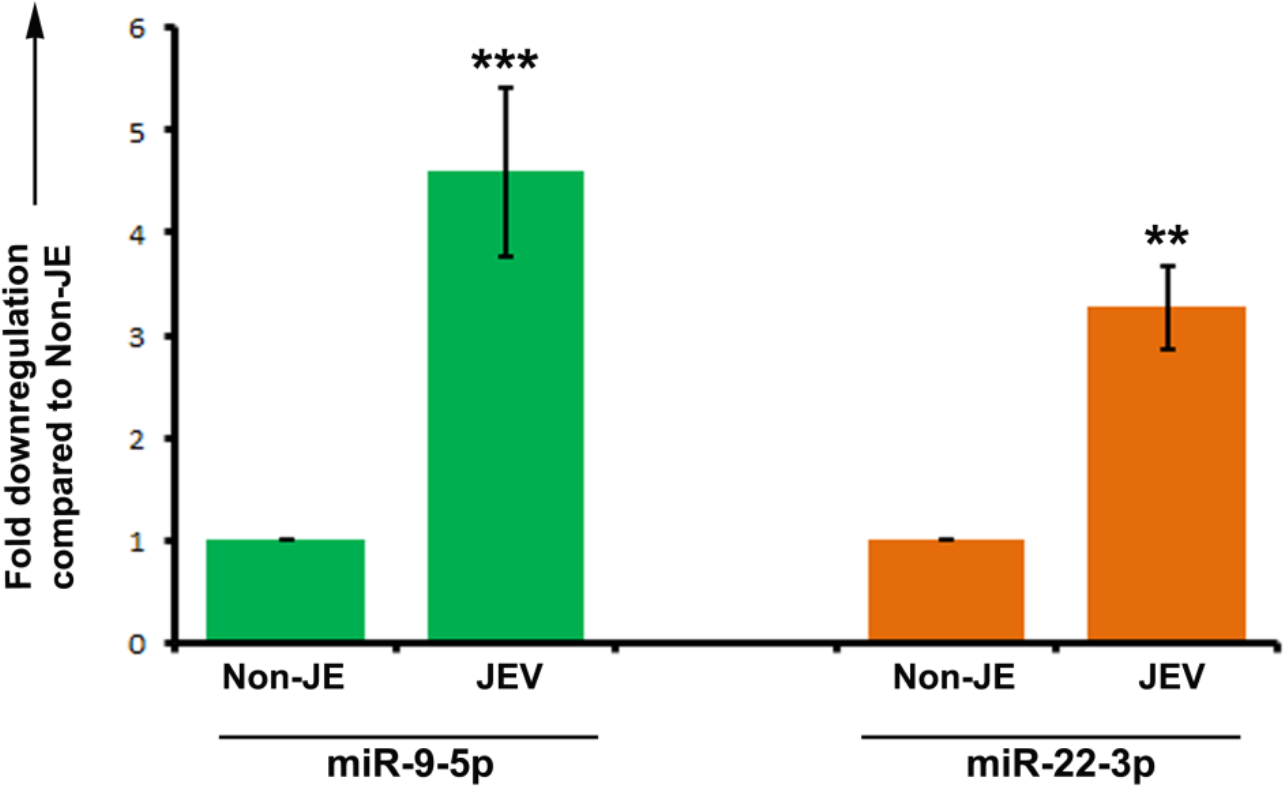
miR-9-5p and miR-22-3p expression in human autopsy tissue of JEV infection. Expression of miR-9-5p and miR-22-3p were analyzed through qRT-PCR in human autopsy tissue of JEV infection and age matched un-infected (non-JE) tissue. Both of the miRNAs were found to be significantly downregulated in virus infected cases as compared to un-infected. Data is representative of three independent experiments (mean± SD) by Student’s t test (**p<0.01, *** p<0.001).

## Discussion

miRNAs play a pivotal role in nervous system cell fate determination in worms^28^, zebra fish^29^ and mammals. miRNAs found in mammalian nervous system is higher than rest of the tissues and the probable reason being its cellular diversity. Same miRNAs can be expressed differently in the different anatomical regions of the brain and again differently during developmental stages. Neurotropic RNA viruses are capable of modulating host miRNA machinery in order to invade and sustain in host system. Changes in human miRNA expression can regulate viral replication, its persistence in host tissue and modulate host immune response as well ^30^. Our research is a pioneer one investigating the miRNA signature of NSPC upon flaviviral infection.

Our data emphasizes on the down regulation of four miRNAs in hNS1 cell line and primary neural precursor cells generated from human foetus. These four miRNAs are extensively reported to play vital role in neuropathology or host-virus interaction. miR-9 is a highly abundant miRNA in vertebrate brain and it affects neurogenesis, neuronal proliferation, differentiation and migration ^31^. miR-9 expression is paramount in sub ventricular zone but it is also expressed in dorsal telencephalon and spinal cord^32,33^. Differential expression of miR-9 is associated with many human neuropathological conditions. In medulloblastoma, reduced miR-9 expression is observed^34^. Contrarily, higher miR-9 expression is seen in glioblastoma^35^. miR-9 also facilitates influenza A infection in A549 cells ^36^. miR-22 is shown to control adult neurogenesis, neuronal migration and dendritic arborization ^37^. It is also considered as a serum biomarker for hepatitis B infection ^38^. miR-124 expression is linked to both pre and post natal neuronal differentiation ^39^.Overexpression of miR-124 suppresses the proliferation of medulloblastoma cells ^40^. miR-132 enhances HIV replication in Jurkat cells^41^. miR-132 is also involved in corneal infection of Herpes simplex virus (HSV)^42^. In case of Influenza A infection in lung cells, miR-132 accumulation is observed^43^. Our report is the first one showing differential expression of these miRNAs in JEV infection.

miRNAs post transcriptionally silent gene expression by degrading the mRNAs. These genes are called miRNA target genes which eventually encode for proteins. We wanted to identify if any protein-protein interaction exists among the miRNA target genes. PPI networks hold a significant importance in system biology. Studying a PPI network provides the intricate details of the biological pathways occurring inside a cell. PPI networks are of immense importance in studying disease biology also. Mutation or differential expression or loss of translation of a protein might affect the complex protein network thus in turn causes functional changes in a biological pathway. A single miRNA controls expression of multiple proteins which among themselves build up a functional network to maintain cellular homeostasis. Therefore, we were interested to evaluate the target gene (protein) interaction of individual miRNAs. The target gene (protein) networks were build up in string database followed by their binarization based on interaction between two genes (proteins). Using community detection algorithms based on iterative greedy optimization methods we identified different modules among gene-gene interactome networks and using hub detection and classification techniques borrowed from graph theory we have further pinpointed key hub genes (namely two classes provincial and connector hub genes) which might play crucial role in the functionality of a given module. High degree genes in the PPI network made significantly increased contributions to structural motif of the module and also tended to have high degree centrality in the interactome network. We separated hub genes into provincial and connector classes, and predicted that the two types of hubs had different functional organization of the remaining network (provincial ones may participate in more within module information processing while the connector class of hub gene may participate in more between module information processing).

To reliably identify hub genes in PPI networks, we used multiple graph theoretical measures, including vertex degree, motif participation coefficient and betweenness and closeness centrality (participation coefficient is being reported here). While hubs are most often identified solely on the basis of their high degree, the relationship of degree to other aspects of their topological embedding is less well understood. While clearly interrelated, each of the measures we apply in this study captures a distinct way in which a community of genes participates in the structure of the whole PPI network.

To test out our prediction about the role of miRNA target gene networks in maintaining the functions of a given module and whether any of the identified genes in a given module showed up-regulated or down-regulated expression post JEV infection we subjected the identified class of hub genes in our study to further analysis in http://www.targetscan.org. Targetsacn is a web server which shows miRNA binding site in the target gene. Out of all the hub genes of four miRNAs we validated the expression of genes which had a binding site for the miRNA as shown in targetscan server. miRNA up-regulation by mimics along with JEV infection emphasizes the key genes which might play important role in viral infection in NSPCs. These were ETS1, IL-6 (miR-9-5p); ESR1 (miR-22-3p); SDC4, PIK3CA, CAV1, IQGAP1 (miR-124-3p). However, miR-132-3p target hub genes have no validated interaction in targetscan, hence being eliminated from validation studies. Notably, all the genes ETS1, IL-6 (miR-9-5p); ESR1 (miR-22-3p); SDC4, PIK3CA, CAV1, IQGAP1 (miR-124-3p) which showed validated interaction in the target scan are connector class of hub genes, and presumably these connector hubs links multiple modules or communities in the PPI network to one another and thus their expression in the cell post viral infection might find critical functional role in JEV pathogenesis as well as maintenance of miRNA machinery.

In summary, we hereby characterize neural stem/progenitor cell specific miRNA profile post JEV infection. These miRNAs might be considered as attractive therapeutic targets against JEV infection. In future we aim to evaluate the intricate molecular regulation of the hub genes of miRNA targets in order to identify their role in flaviviral infection.

## Acknowledgement

This project is funded by Department of Biotechnology, Government of India research grant to AB (Anirban Basu, Grant No.BT/PR22341/MED/122/55/2016). AB (Anirban Basu) is also a recipient of Tata Innovation Fellowship (BT/HRD/35/01/02/2014) from the Department of Biotechnology. DR is supported by the Department of Biotechnology, Government of India, Ramalingaswami fellowship (BT/RLF/Re-entry/07/2014) and DST-CSRI Government of India extramural research grant (SR/CSRI/21/2016) and NBRC core funding. SM is a recipient of DST-INSPIRE fellowship, Government of India (IF140074). We sincerely acknowledge Prof. Anita Mahadevan and Prof. SK Shankar, Department of Neuropathology, National Institute for Mental Health and Neurosciences, Bangalore, for providing uninfected and JE infected autopsied human tissue. Distributed Information Centre (DIC) of NBRC is acknowledged for computer related infrastructural support. Additionally, we acknowledge Kanhaiya Kumawat and Manish Dogra for their technical help.

## References

1 Mukherjee, S. et al. Japanese encephalitis virus induces human neural stem/progenitor cell death by elevating GRP78, PHB and hnRNPC through ER stress. Cell death & disease 8, e2556, doi:10.1038/cddis.2016.394 (2017).

2 Das, S. & Basu, A. Japanese encephalitis virus infects neural progenitor cells and decreases their proliferation. Journal of neurochemistry 106, 1624–1636, doi:10.1111/j.1471-4159.2008.05511.x (2008).

3 Liu, Q., Zhang, L. & Li, H. New Insights: MicroRNA Function in CNS Development and Psychiatric Diseases. Current Pharmacology Reports 4, 132–144, doi:10.1007/s40495-018-0129-2 (2018).

4 Shi, Y. et al. MicroRNA regulation of neural stem cells and neurogenesis. The Journal of neuroscience: the official journal of the Society for Neuroscience 30, 14931–14936, doi:10.1523/JNEUROSCI.4280-10.2010 (2010).

5 Lee, R. C., Feinbaum, R. L. & Ambros, V. The C. elegans heterochronic gene lin-4 encodes small RNAs with antisense complementarity to lin-14. Cell 75, 843–854 (1993).

6 Gangaraju, V. K. & Lin, H. MicroRNAs: key regulators of stem cells. Nature reviews. Molecular cell biology 10, 116–125, doi:10.1038/nrm2621 (2009).

7 Mathieu, J. & Ruohola-Baker, H. Regulation of stem cell populations by microRNAs. Advances in experimental medicine and biology 786, 329–351, doi:10.1007/978-94-007-6621-1_18 (2013).

8 Femminella, G. D., Ferrara, N. & Rengo, G. The emerging role of microRNAs in Alzheimer’s disease. Frontiers in physiology 6, 40, doi:10.3389/fphys.2015.00040 (2015).

9 Leggio, L. et al. microRNAs in Parkinson’s Disease: From Pathogenesis to Novel Diagnostic and Therapeutic Approaches. International journal of molecular sciences 18, doi:10.3390/ijms18122698 (2017).

10 Hoye, M. L. et al. MicroRNA Profiling Reveals Marker of Motor Neuron Disease in ALS Models. The Journal of neuroscience: the official journal of the Society for Neuroscience 37, 5574–5586, doi:10.1523/JNEUROSCI.3582-16.2017 (2017).

11 Caputo, V., Ciolfi, A., Macri, S. & Pizzuti, A. The emerging role of MicroRNA in schizophrenia. CNS & neurological disorders drug targets 14, 208–221 (2015).

12 Hicks, S. D. & Middleton, F. A. A Comparative Review of microRNA Expression Patterns in Autism Spectrum Disorder. Frontiers in psychiatry 7, 176, doi:10.3389/fpsyt.2016.00176 (2016).

13 Simonson, B. & Das, S. MicroRNA Therapeutics: the Next Magic Bullet? Mini reviews in medicinal chemistry 15, 467–474 (2015).

14 Ho, B. C. et al. Inhibition of miR-146a prevents enterovirus-induced death by restoring the production of type I interferon. Nature communications 5, 3344, doi:10.1038/ncomms4344 (2014).

15 Wu, S. et al. miR-146a facilitates replication of dengue virus by dampening interferon induction by targeting TRAF6. The Journal of infection 67, 329–341, doi:10.1016/j.jinf.2013.05.003 (2013).

16 Trobaugh, D. W. et al. RNA viruses can hijack vertebrate microRNAs to suppress innate immunity. Nature 506, 245–248, doi:10.1038/nature12869 (2014).

17 Song, L., Liu, H., Gao, S., Jiang, W. & Huang, W. Cellular microRNAs inhibit replication of the H1N1 influenza A virus in infected cells. Journal of virology 84, 8849–8860, doi:10.1128/JVI.00456-10 (2010).

18 Zheng, Z. et al. Human microRNA hsa-miR-296-5p suppresses enterovirus 71 replication by targeting the viral genome. Journal of virology 87, 5645–5656, doi:10.1128/JVI.02655-12 (2013).

19 Fatima, M. et al. Tripartite containing motif 32 modulates proliferation of human neural precursor cells in HIV-1 neurodegeneration. Cell death and differentiation 23, 776–786, doi:10.1038/cdd.2015.138 (2016).

20 Zhu, B. et al. MicroRNA-15b Modulates Japanese Encephalitis Virus-Mediated Inflammation via Targeting RNF125. Journal of immunology 195, 2251–2262, doi:10.4049/jimmunol.1500370 (2015).

21 Thounaojam, M. C., Kaushik, D. K., Kundu, K. & Basu, A. MicroRNA-29b modulates Japanese encephalitis virus-induced microglia activation by targeting tumor necrosis factor alpha-induced protein 3. Journal of neurochemistry 129, 143–154, doi:10.1111/jnc.12609 (2014).

22 Thounaojam, M. C. et al. MicroRNA 155 regulates Japanese encephalitis virus-induced inflammatory response by targeting Src homology 2-containing inositol phosphatase 1. Journal of virology 88, 4798–4810, doi:10.1128/JVI.02979-13 (2014).

23 Hazra, B., Kumawat, K. L. & Basu, A. The host microRNA miR-301a blocks the IRF1-mediated neuronal innate immune response to Japanese encephalitis virus infection. Science signaling 10, eaaf5185, doi:10.1126/scisignal.aaf5185 (2017).

24 Mukherjee, S. et al. PLVAP and GKN3 Are Two Critical Host Cell Receptors Which Facilitate Japanese Encephalitis Virus Entry Into Neurons. Scientific reports 8, 11784, doi:10.1038/s41598-018-30054-z (2018).

25 Rubinov, M. & Sporns, O. Complex network measures of brain connectivity: uses and interpretations. NeuroImage 52, 1059–1069, doi:10.1016/j.neuroimage.2009.10.003 (2010).

26 Newman, M. E. Modularity and community structure in networks. Proceedings of the National Academy of Sciences of the United States of America 103, 8577–8582, doi:10.1073/pnas.0601602103 (2006).

27 Sporns, O., Honey, C. J. & Kotter, R. Identification and classification of hubs in brain networks. PloS one 2, e1049, doi:10.1371/journal.pone.0001049 (2007).

28 Johnston, R. J. & Hobert, O. A microRNA controlling left/right neuronal asymmetry in Caenorhabditis elegans. Nature 426, 845–849, doi:10.1038/nature02255 (2003).

29 Giraldez, A. J. et al. MicroRNAs regulate brain morphogenesis in zebrafish. Science 308, 833–838, doi:10.1126/science.1109020 (2005).

30 Umbach, J. L. & Cullen, B. R. The role of RNAi and microRNAs in animal virus replication and antiviral immunity. Genes & development 23, 1151–1164, doi:10.1101/gad.1793309 (2009).

31 Radhakrishnan, B. & Alwin Prem Anand, A. Role of miRNA-9 in Brain Development. Journal of experimental neuroscience 10, 101–120, doi:10.4137/JEN.S32843 (2016).

32 Otaegi, G., Pollock, A., Hong, J. & Sun, T. MicroRNA miR-9 modifies motor neuron columns by a tuning regulation of FoxP1 levels in developing spinal cords. The Journal of neuroscience: the official journal of the Society for Neuroscience 31, 809–818, doi:10.1523/JNEUROSCI.4330-10.2011 (2011).

33 Leucht, C. et al. MicroRNA-9 directs late organizer activity of the midbrain-hindbrain boundary. Nature neuroscience 11, 641–648, doi:10.1038/nn.2115 (2008).

34 Ferretti, E. et al. MicroRNA profiling in human medulloblastoma. International journal of cancer 124, 568–577, doi:10.1002/ijc.23948 (2009).

35 Kim, T. M., Huang, W., Park, R., Park, P. J. & Johnson, M. D. A developmental taxonomy of glioblastoma defined and maintained by MicroRNAs. Cancer research 71, 3387–3399, doi:10.1158/0008-5472.CAN-10-4117 (2011).

36 Dong, C., Sun, X., Guan, Z., Zhang, M. & Duan, M. Modulation of influenza A virus replication by microRNA-9 through targeting MCPIP1. Journal of medical virology 89, 41–48, doi:10.1002/jmv.24604 (2017).

37 Beamer, E. H. et al. MicroRNA-22 Controls Aberrant Neurogenesis and Changes in Neuronal Morphology After Status Epilepticus. Frontiers in molecular neuroscience 11, 442, doi:10.3389/fnmol.2018.00442 (2018).

38 Sarkar, N. & Chakravarty, R. Hepatitis B Virus Infection, MicroRNAs and Liver Disease. International journal of molecular sciences 16, 17746–17762, doi:10.3390/ijms160817746 (2015).

39 Smirnova, L. et al. Regulation of miRNA expression during neural cell specification. The European journal of neuroscience 21, 1469–1477, doi:10.1111/j.1460-9568.2005.03978.x (2005).

40 Pierson, J., Hostager, B., Fan, R. & Vibhakar, R. Regulation of cyclin dependent kinase 6 by microRNA 124 in medulloblastoma. Journal of neuro-oncology 90, 1–7, doi:10.1007/s11060-008-9624-3 (2008).

41 Chiang, K., Liu, H. & Rice, A. P. miR-132 enhances HIV-1 replication. Virology 438, 1–4, doi:10.1016/j.virol.2012.12.016 (2013).

42 Mulik, S. et al. Role of miR-132 in angiogenesis after ocular infection with herpes simplex virus. The American journal of pathology 181, 525–534, doi:10.1016/j.ajpath.2012.04.014 (2012).

43 Buggele, W. A., Johnson, K. E. & Horvath, C. M. Influenza A virus infection of human respiratory cells induces primary microRNA expression. The Journal of biological chemistry 287, 31027–31040, doi:10.1074/jbc.M112.387670 (2012).

